## Supplementary figures and information for Moniruzzaman et al, 2019. for "Dynamic Genome Evolution and Blueprint of Complex Virocell Metabolism in Globally-Distributed Giant Viruses"

### Supplementary datasets and figures for Moniruzzaman et al., 2019.

#### Supplementary dataset descriptions

**Dataset S1.** This spreadsheet contains summary information for each NCLDV MAG reported in this study, including the genome size, number of encoded proteins, metagenome of origin, and other data. Additionally, this spreadsheet contains contig-level summary statistics for all decontamination analyses used in this study, and a list of all contigs that were removed due to similarity to cellular or bacteriophage sources. Lastly, this sheet contains a list of all reference NCLDV genomes used in this study.

**Dataset S2.** This spreadsheet contains the annotation information for the orthologous groups calculated in this study. Only orthologous groups that exhibited similarity to a known protein family are listed here. Also provided are the enriched COG categories identified in each NCLDV clade.

**These datasets are available on the Aylward Lab Figshare account:**

https://figshare.com/authors/Frank_Aylward/5008391 in the project titled “NCLDV”.

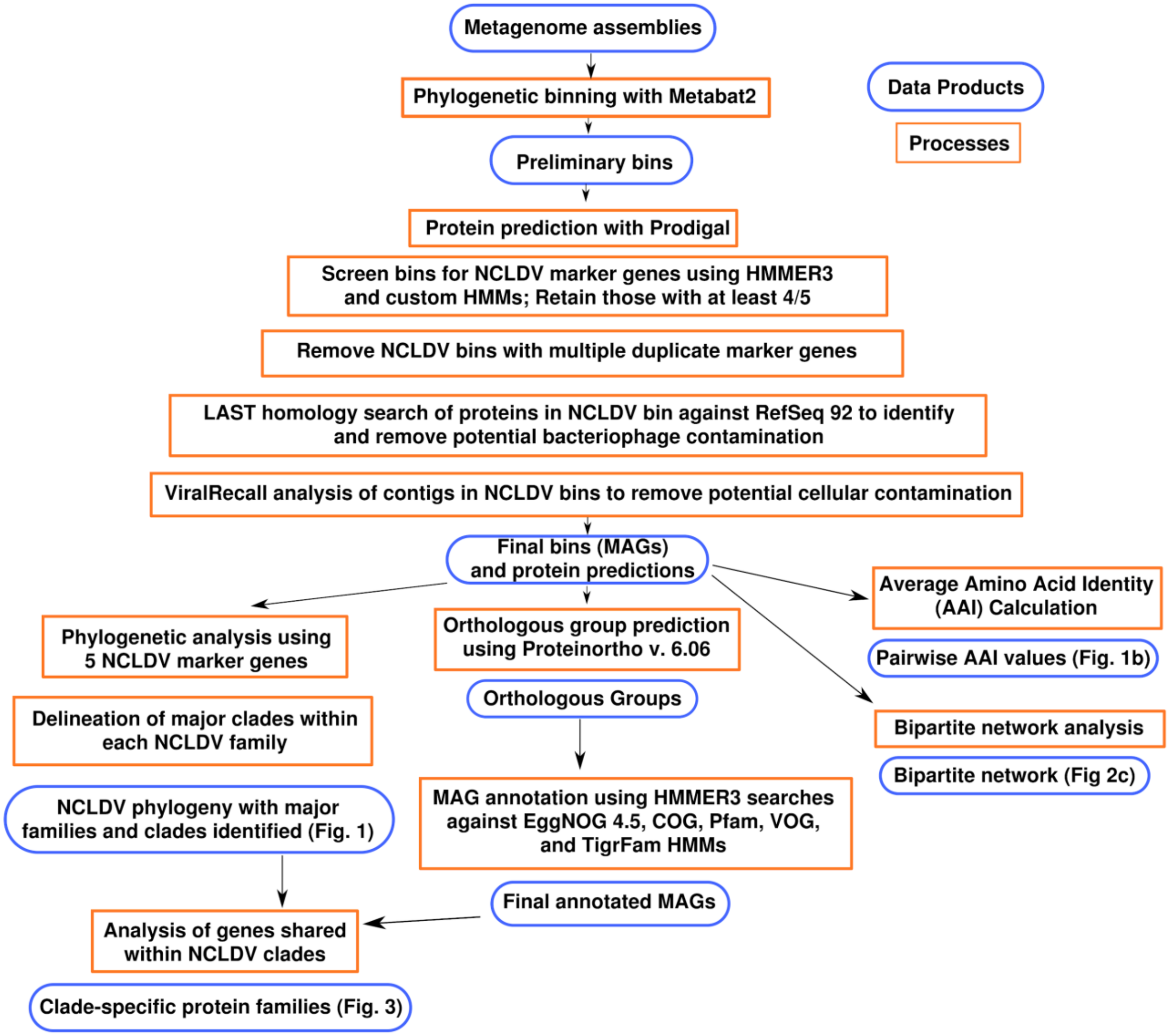


**Figure S1.** Overview of the workflow used to identify and analyze the NCLDV MAGs described in this study.

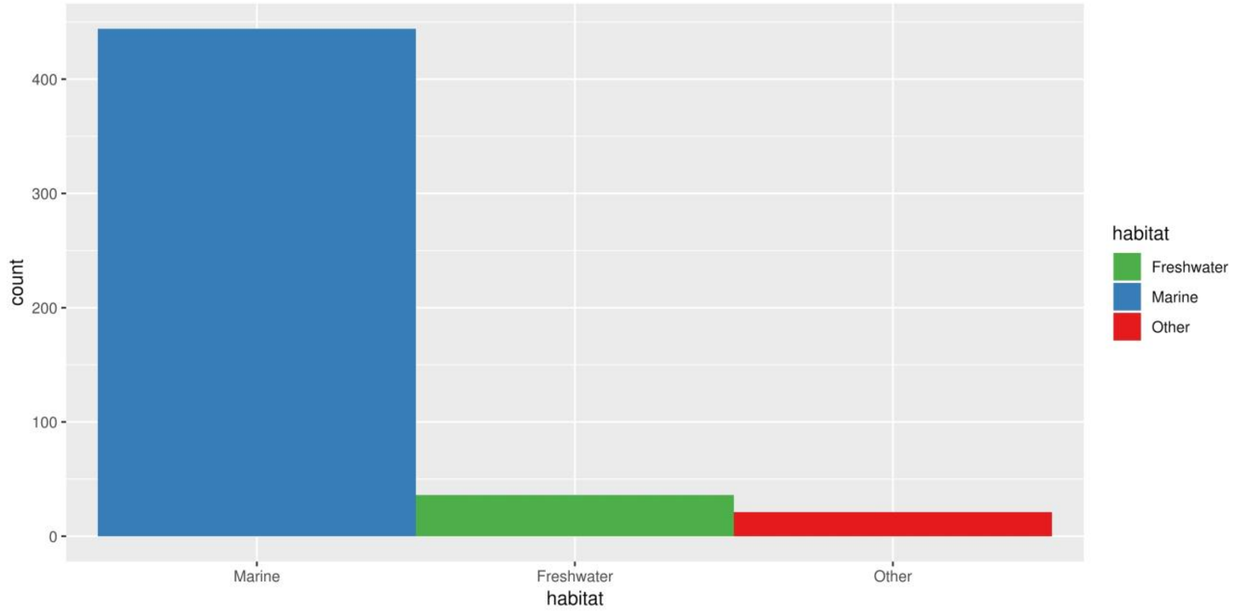


**Figure S2.** Distribution of the habitats represented by the 501 NCLDV MAGs presented in this study.

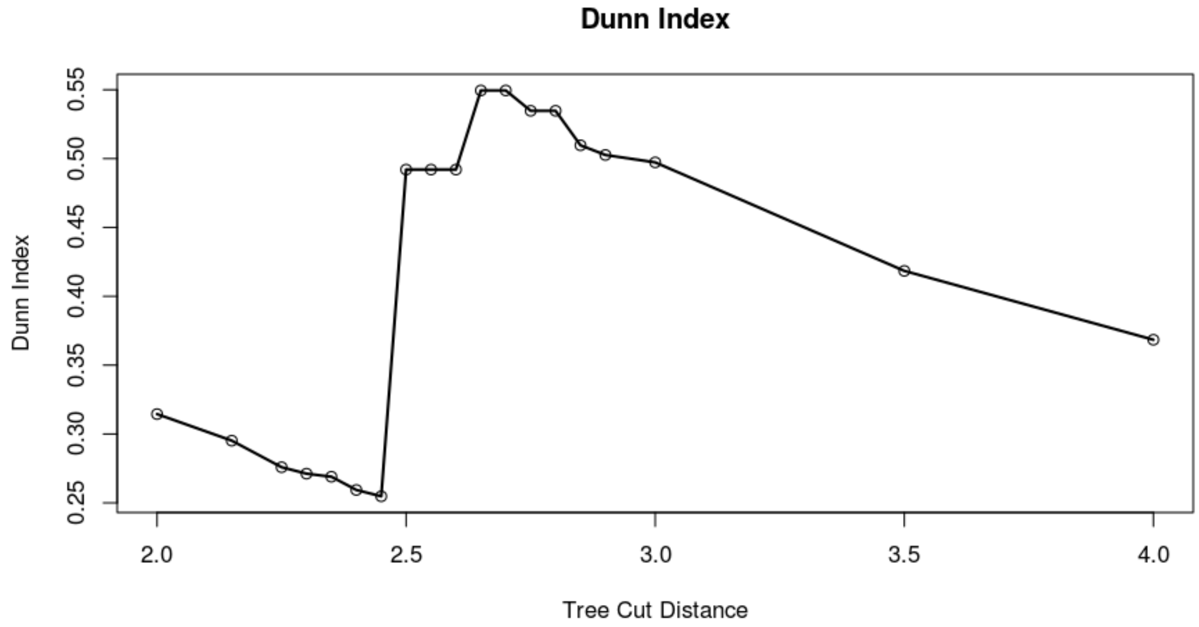


**Figure S3.** Dunn index values for the different tree cut heights in our multi-locus NCLDV phylogeny. A cut height of 2.45 provided the best quality clade-level delineations, and these clusters were subsequently manually edited and inspected to provide the final NCLDV clades.

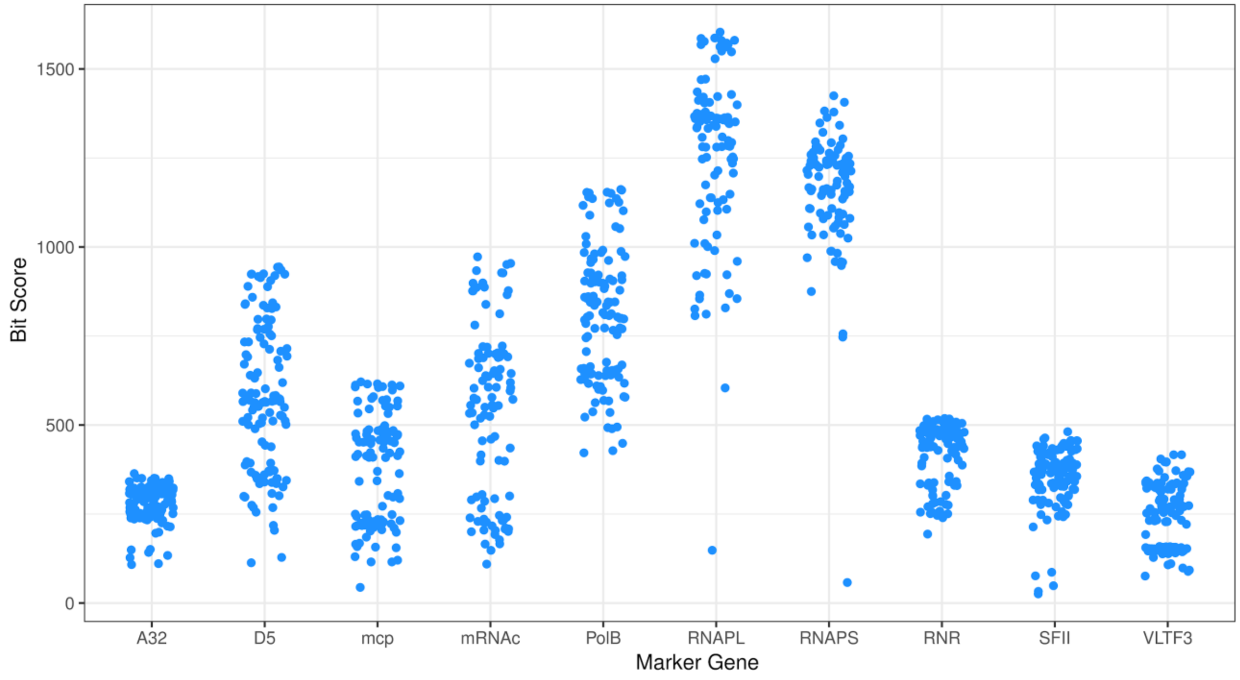


**Figure S4.** Benchmarking of the Megavirales marker gene HMMs designed in this study. The bit scores are for the best hits recovered for each of 121 reference Megavirales genomes in NCBI GenBank. These HMMs were subsequently used to identify marker genes in the Megavirales MAGs generated in this study. The genes A32, MCP, PolB, SFII, and VLTF3 were subsequently used for phylogenetic analyses given they are broadly found across all Megavirales families.

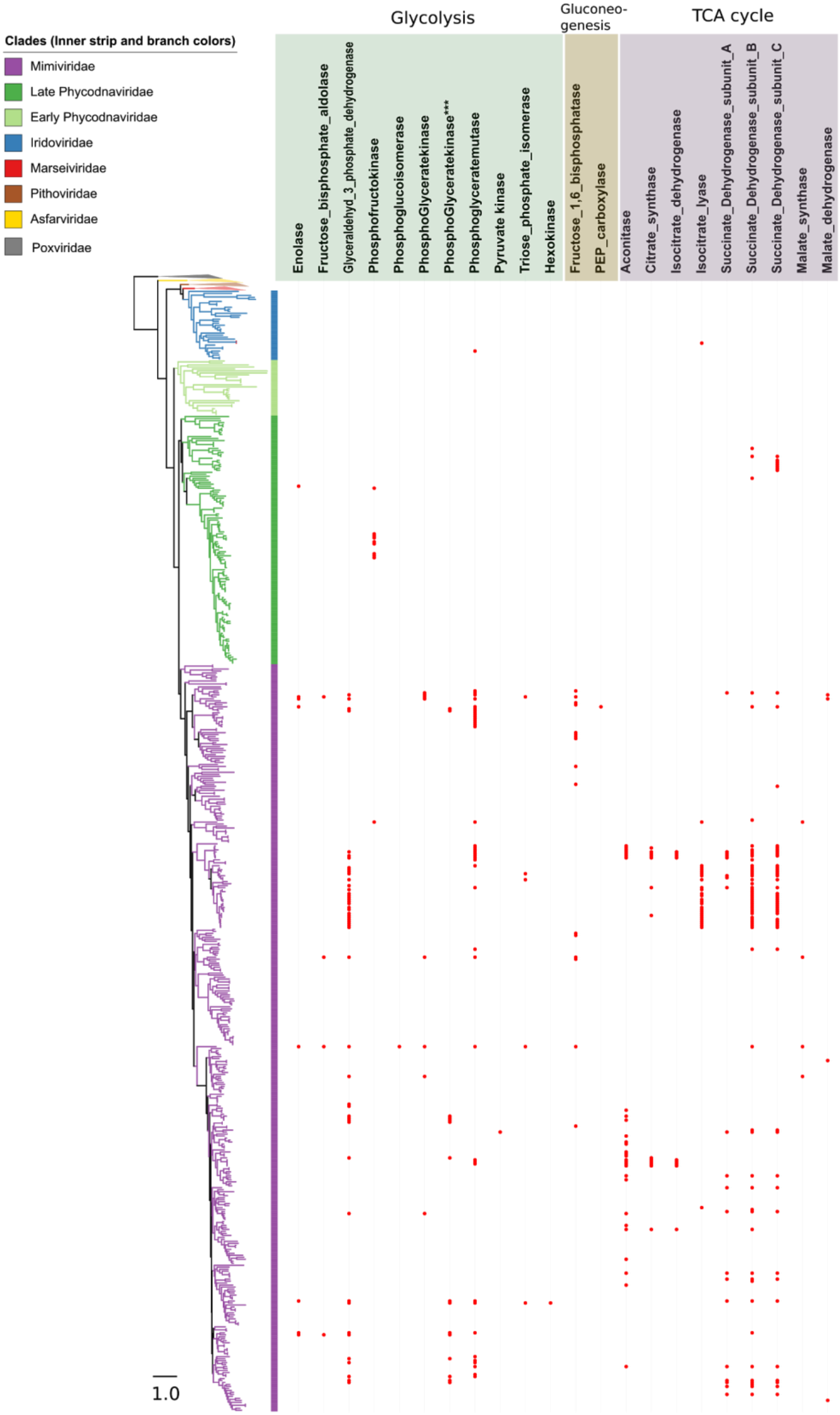


**Figure S5.** Presence/absence of genes involved in Glycolysis, gluconeogenesis and TCA cycle in the NCLDV genomes. Pox-, Pitho- and Asfarviridae families were collapsed since none of these genes were found in the studied genomes from these families.

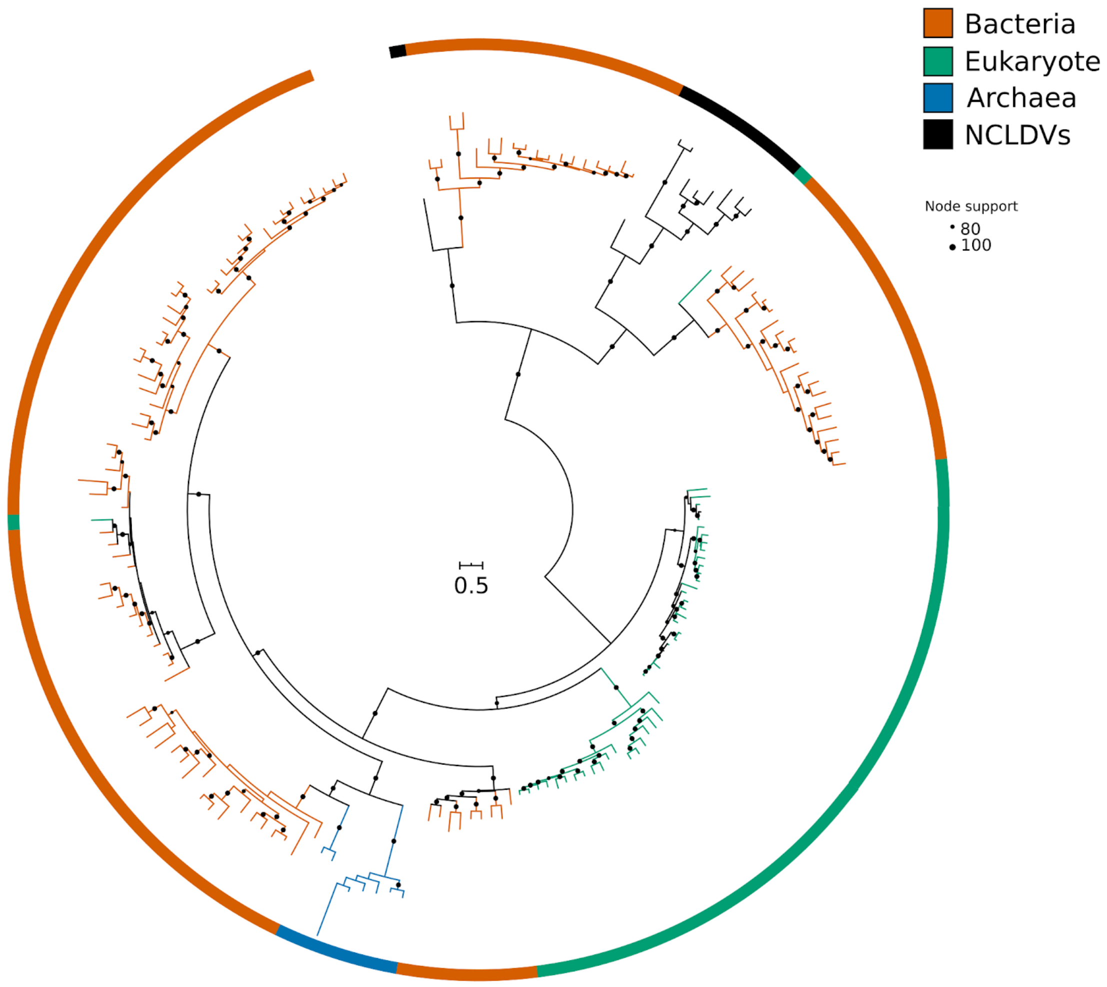


**Figure S6 (A):** Maximum-likelihood phylogenetic reconstruction of glycolytic enzyme phosphofructokinase. Ultrafast bootstrap support >80% are shown.

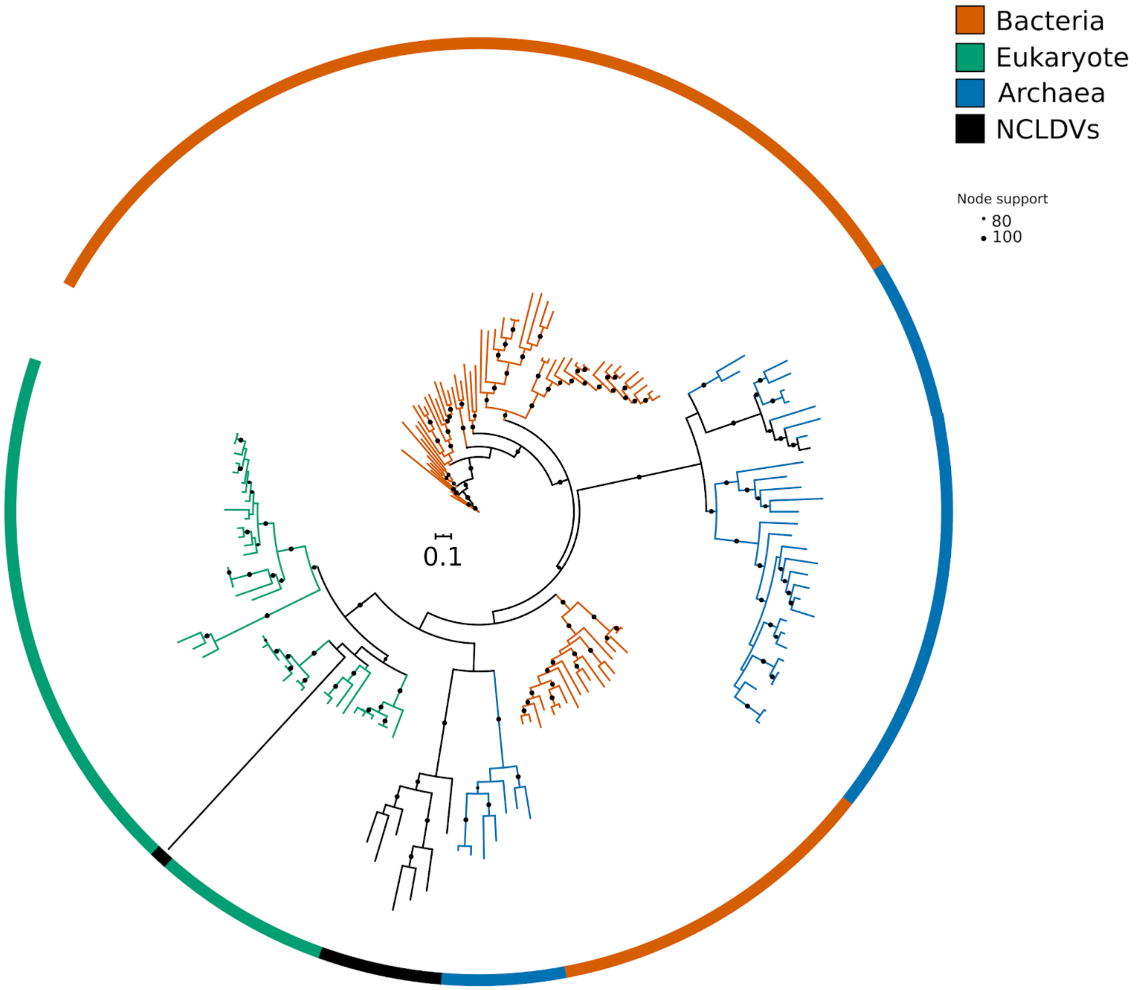


**Figure S6 (B):** Maximum-likelihood phylogenetic reconstruction of the glycolytic enzyme phosphopyruvate hydratase (enolase). Ultrafast bootstrap support >80% are shown.

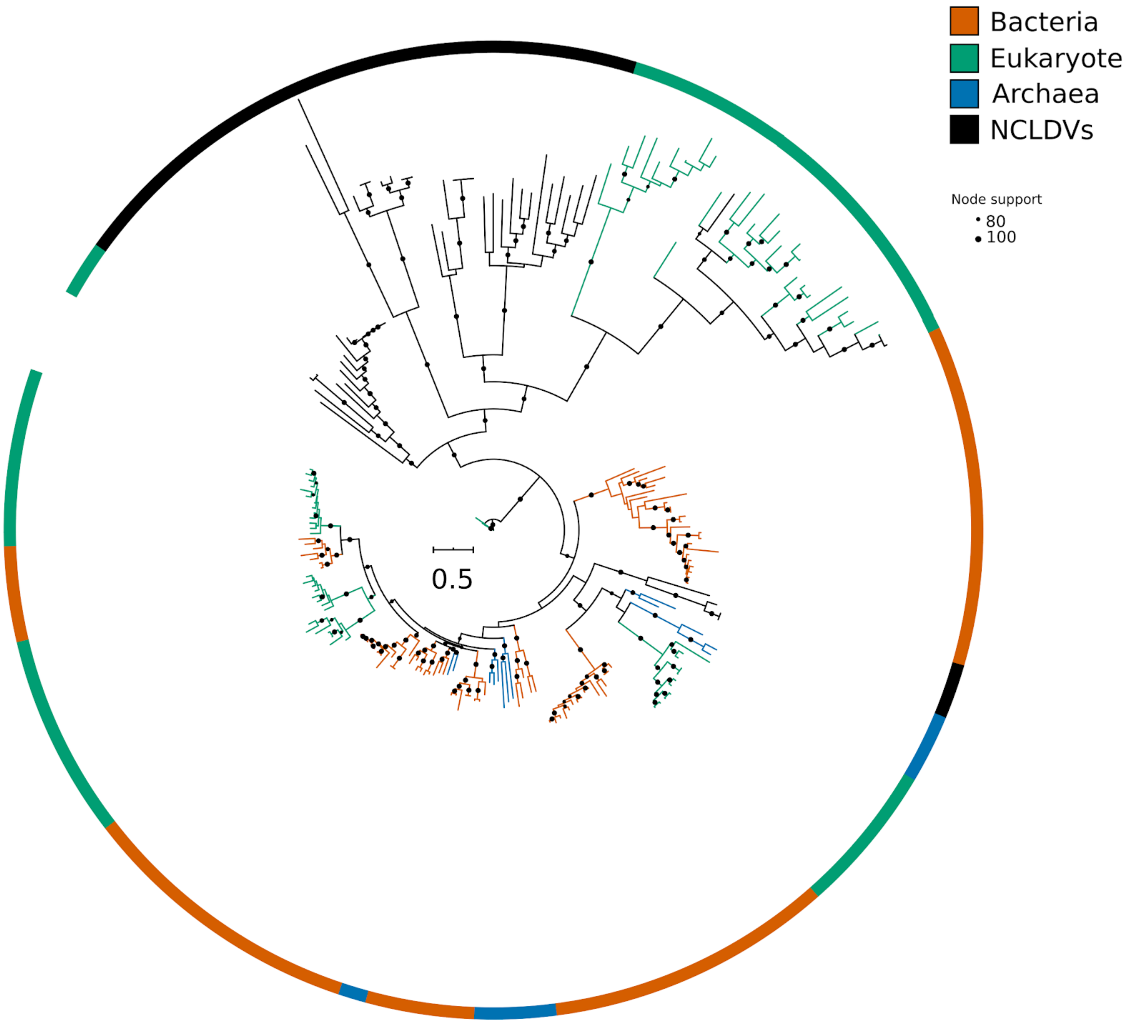


**Figure S6 (C) :** Maximum-likelihood phylogenetic reconstruction of the glycolytic enzyme phosphoglycerate mutase. Ultrafast bootstrap support >80% are shown.

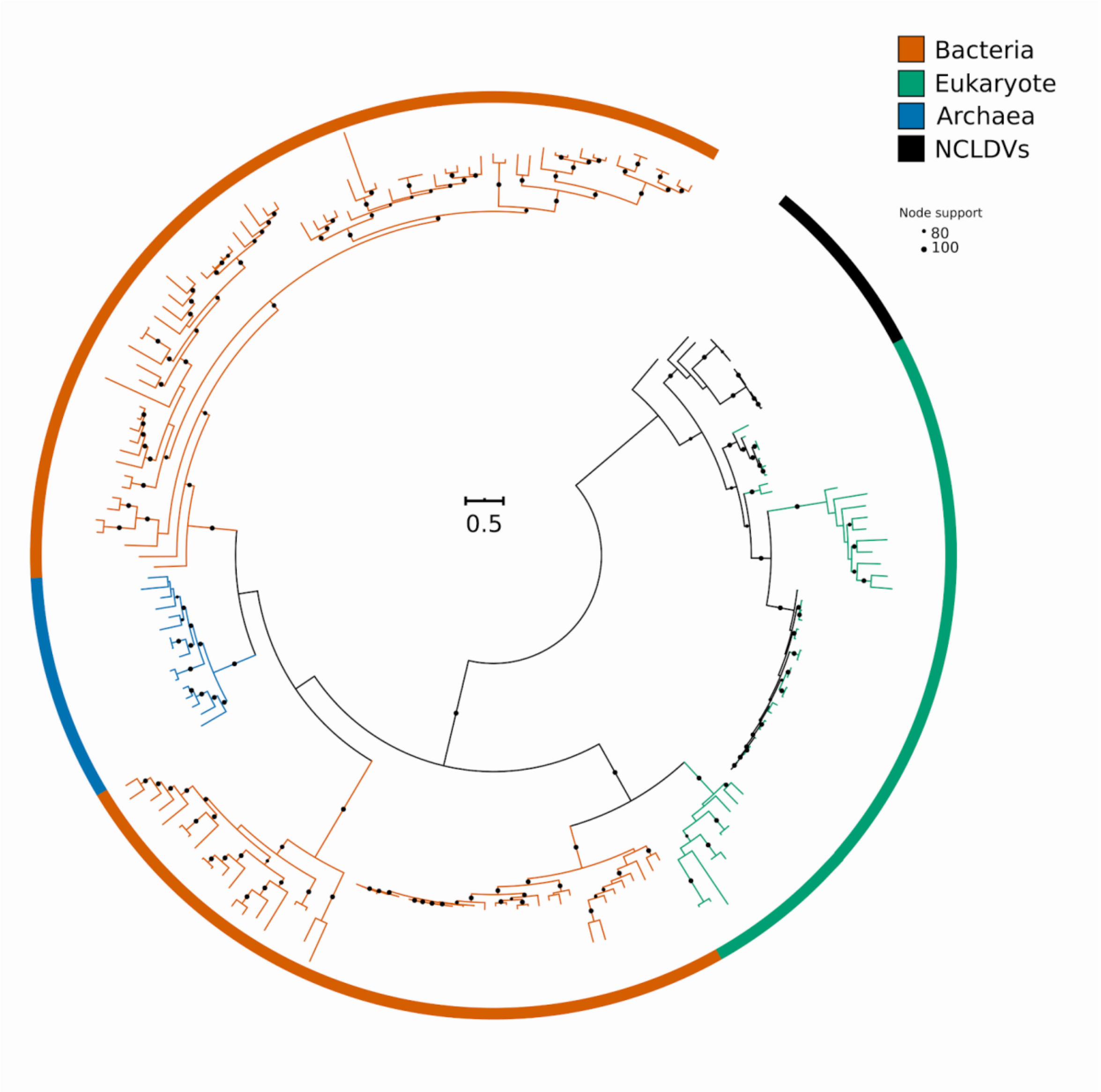


**Figure S6 (D):** Maximum-likelihood phylogenetic reconstruction of the TCA cycle enzyme citrate synthase. Ultrafast bootstrap support >80% are shown.

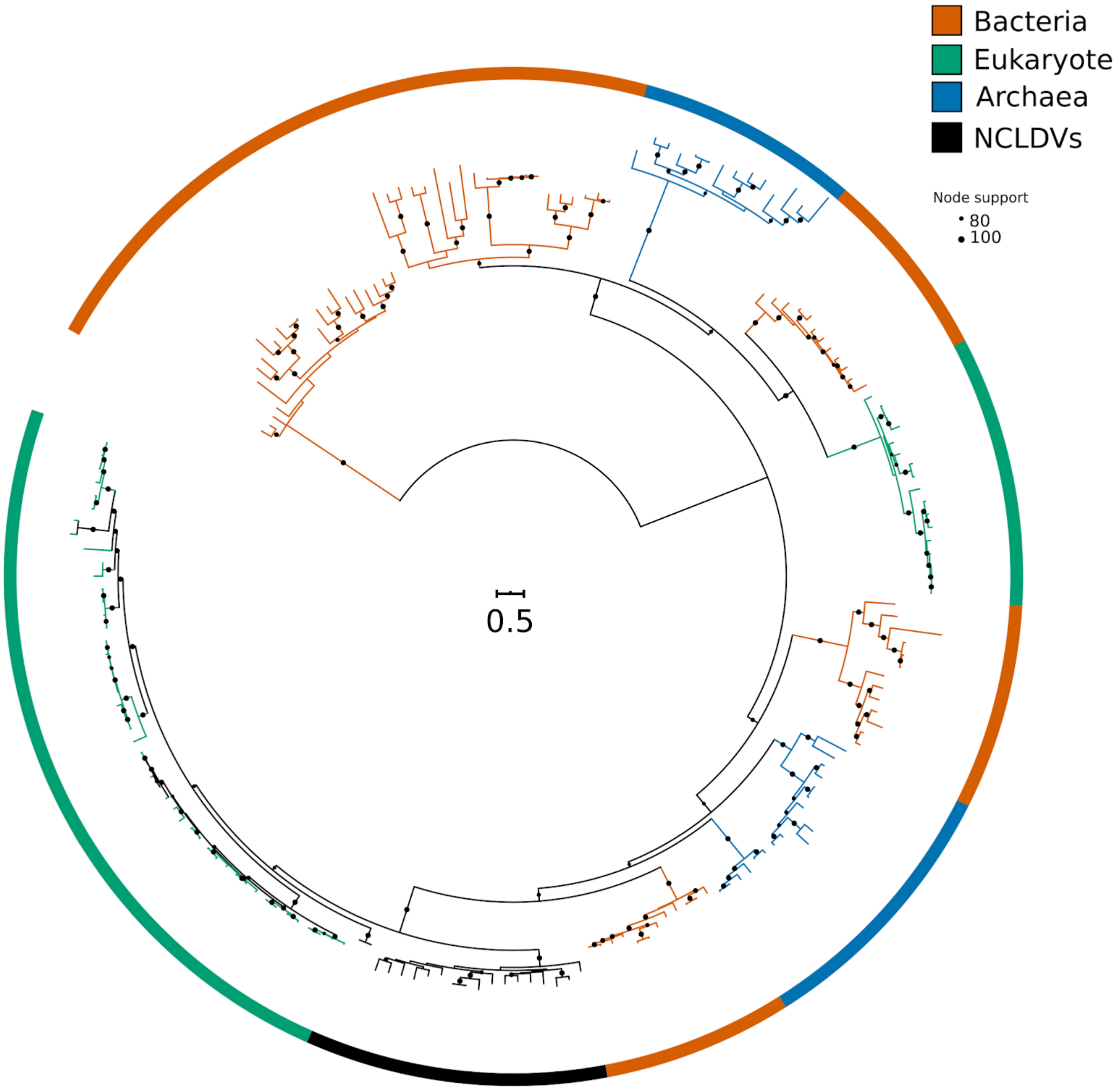


**Figure S6 (E):** Maximum-likelihood phylogenetic reconstruction of the TCA cycle enzyme Succinate dehydrogenase subunit A. Ultrafast bootstrap support >80% are shown.

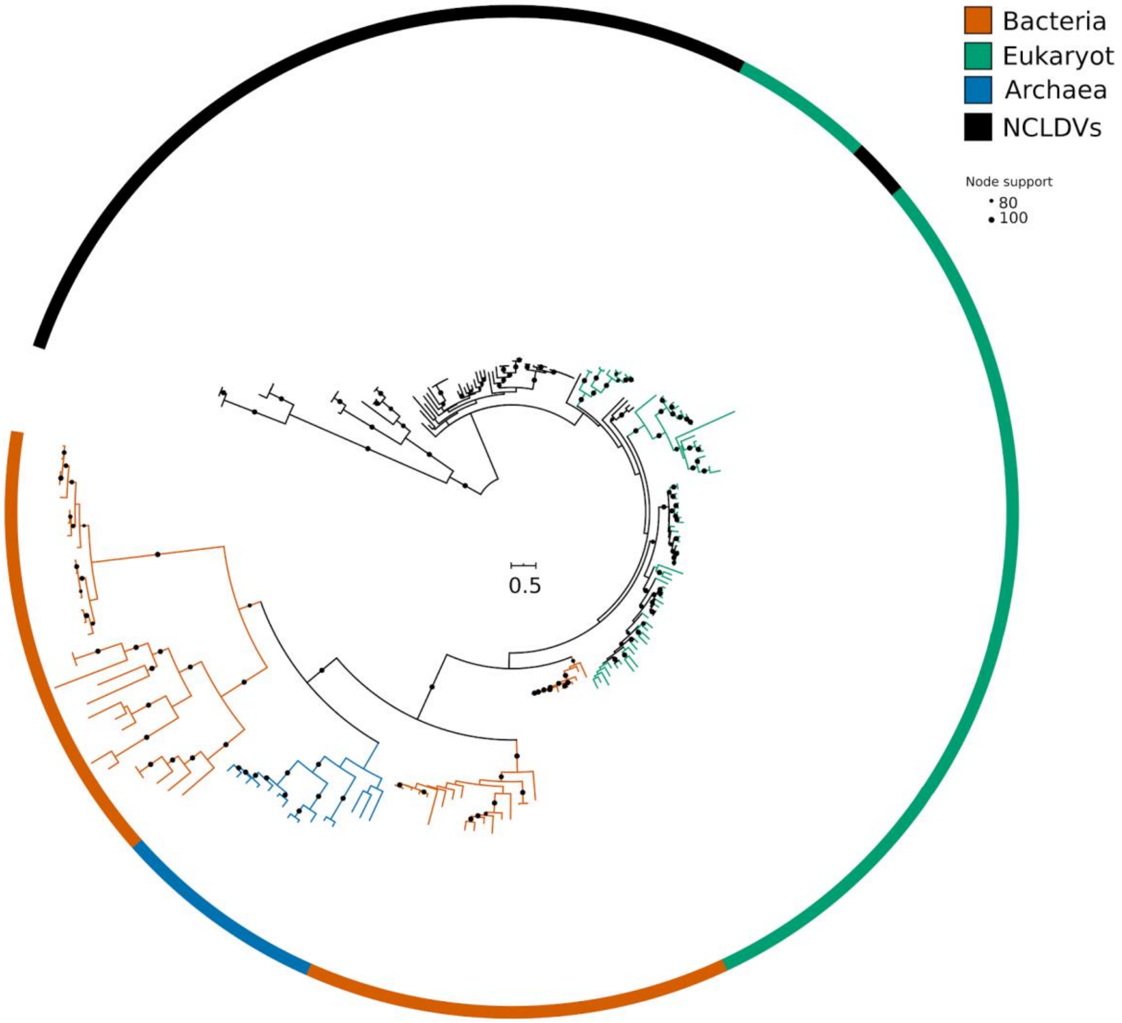


**Figure S6 (F):** Maximum-likelihood phylogenetic reconstruction of the TCA cycle enzyme Succinate dehydrogenase subunit B. Ultrafast bootstrap support >80% are shown.

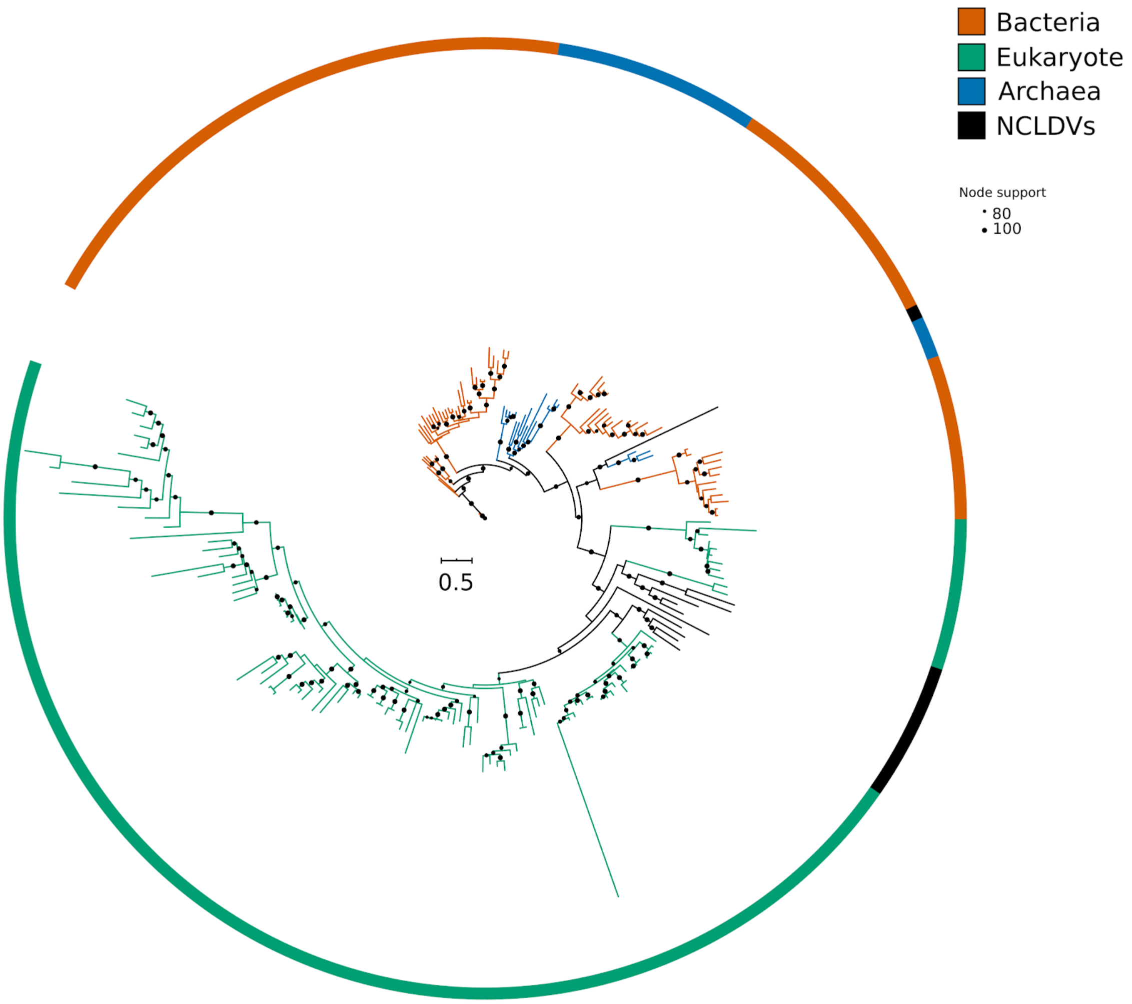


193
194 **Figure S6 (G):** Maximum-likelihood phylogenetic reconstruction of the iron storage and
195 homoeostasis protein ferritin. Ultrafast bootstrap support >80% are shown.
196

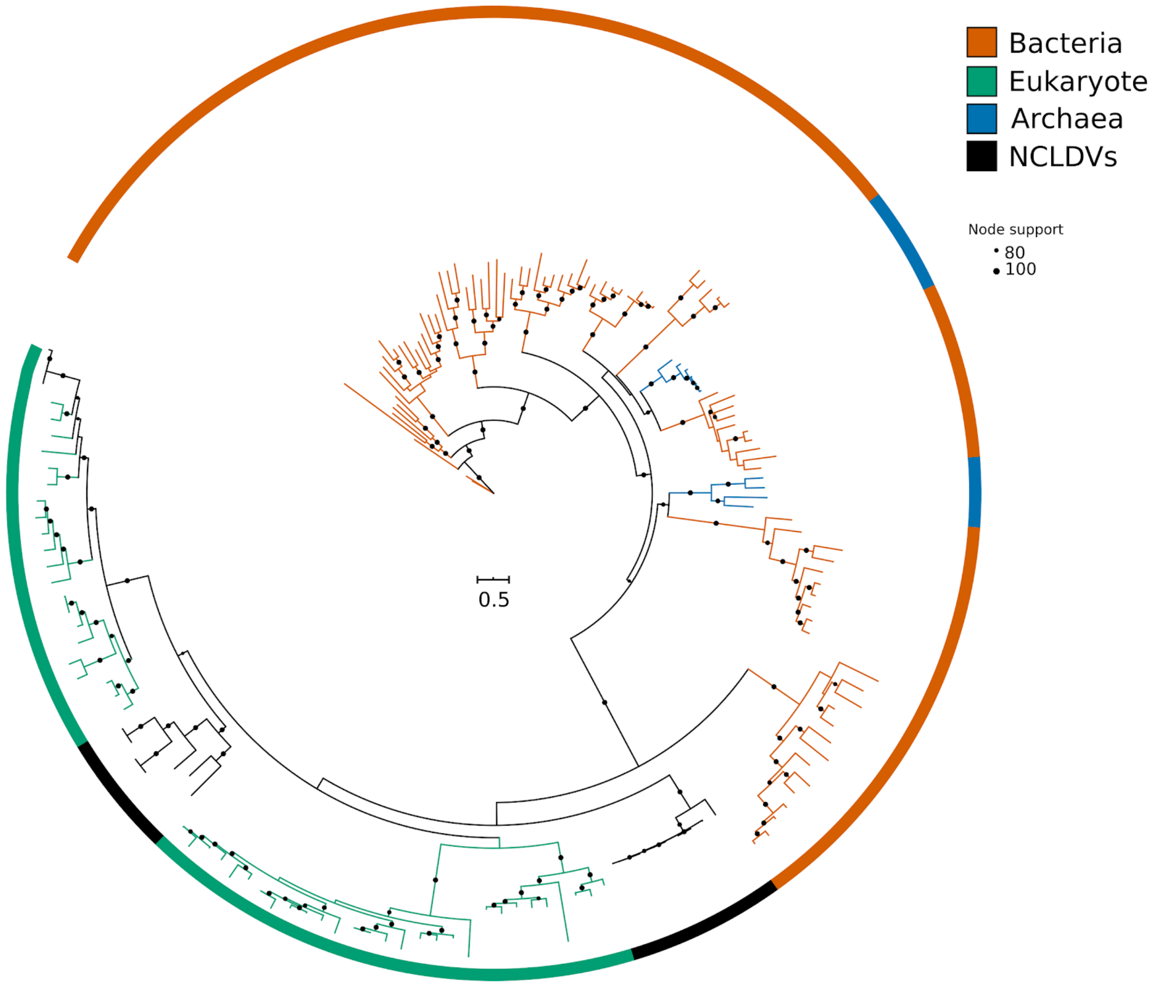


**Figure S6 (H):** Maximum-likelihood phylogenetic reconstruction of the Phosphate/Na+ symporter family proteins. Ultrafast bootstrap support >80% are shown.

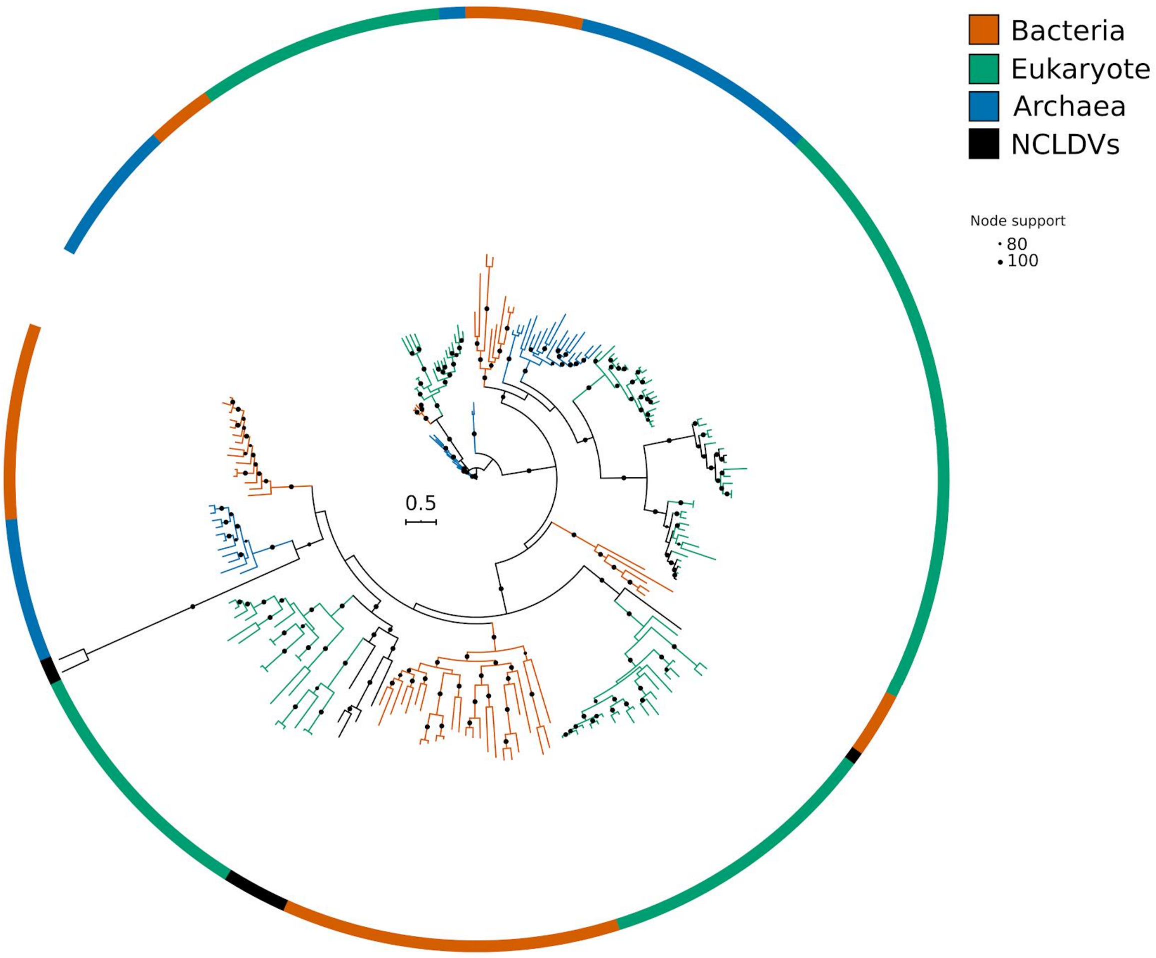


**Figure S6 (I):** Maximum-likelihood phylogenetic reconstruction of the Ammonium transporter (AmT) proteins. Ultrafast bootstrap support >80% are shown.

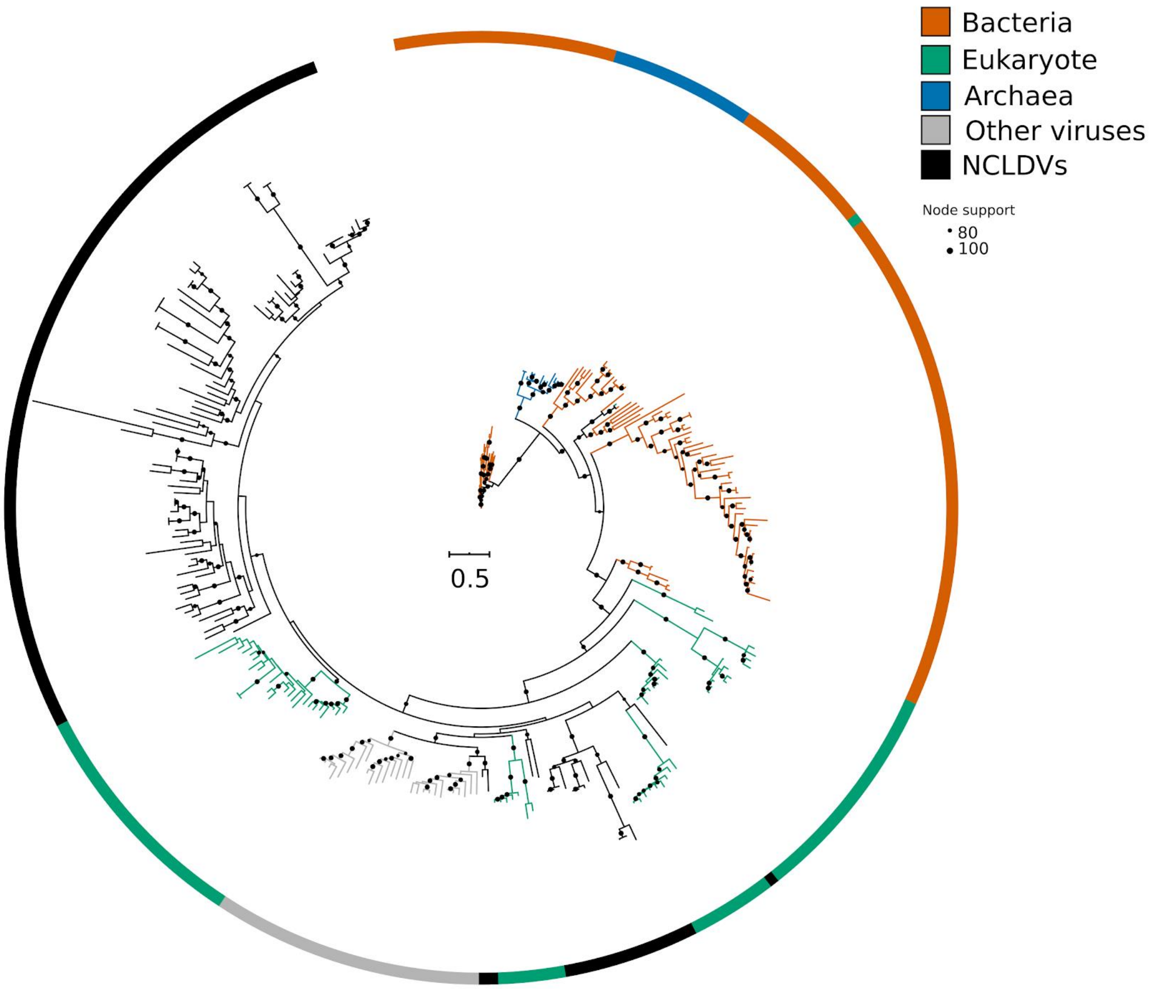


**Figure S6 (J):** Maximum-likelihood phylogenetic reconstruction of the antioxidant enzyme superoxide dismutase (SOD). Ultrafast bootstrap support >80% are shown. “Other viruses” refer to sequences from eukaryotic viruses other than NCLDVs.
